# The size of the functional base of support decreases with age

**DOI:** 10.1101/2025.05.19.654897

**Authors:** Lizeth H. Sloot, Thomas Gerhardy, Katja Mombaur, Matthew Millard

## Abstract

Falls occur more often as we age. To identify people at risk of falling, balance analysis requires an accurate base-of-support model. We previously developed a functional base-of-support (fBOS) model for standing young adults and showed that its area is smaller than the footprint area. Our fBOS model is a polygon that contains centre-of-pressure (COP) trajectories recorded as standing participants move their COP in the largest possible loop while keeping their feet flat on the ground. Here we assess how the size of the fBOS changes with age by comparing 38 younger (YA), 14 middle-aged (MA), and 34 older adults (OA). The fBOS area is smaller in older adults: OA area is 58% of the YA area (*p* < 0.001), and 59% of the MA area (*p* = 0.001), with no difference between YA and MA. The reduction in fBOS area among the OA is primarily caused by a reduction in the length of the fBOS. In addition, among older adults smaller fBOS areas correlated with a lower score on the Short Physical Performance Battery (*τ*=0.28, *p* = 0.04), a reduced walking speed (*τ*=0.25, *p* = 0.04), and a higher frailty level (*p* = 0.09). So that others can extend our work, we have made our fBOS models available online.

## Introduction

Falls are a recurrent problem in the ageing population, with one in every three adults above 65 years of age falling yearly^1–3^. Analysing balance is challenging because it is affected by the three-dimensional movements of all of the body’s segments and is particularly influenced by foot placement location. Models of dynamic balance, such as the extrapolated center of mass^4^ and foot placement estimator^5–7^, simplify balance analysis by using the position and velocity of the body’s segments to calculate the location of a balance point on the floor: if the BOS covers this point the body is considered to be balanced according to the assumptions of the model. The distance between the balance point and the nearest BOS edge (called the margin-of-stability) is often analysed because it provides a measure of how close someone’s body is to the boundary that separates being balanced from being off-balance. The accuracy of the margin-of-stability depends on both the balance model and the BOS model.

Unfortunately, there is no standardised definition of the geometry of the BOS. Many balance studies rely on an ad-hoc definition for the nearest BOS edge when evaluating the margin-of-stability, often using either just a single toe or heel motion capture marker^8^. This approach assumes that the BOS is the same as the footprint, but evidence shows that the area that can functionally support a large fraction of the body’s weight is much smaller^9,10^. Our previous work^10^ suggests that using the footprint as the BOS can introduce 2.3-5.0 cm of error into the margin-of-stability for an average-sized adult foot. Although this may seem small, it is large in comparison to the dimensions of the fBOS and can mean the difference between remaining stably balanced and needing to take a compensatory step.

While the fBOS has been measured in a number of studies, the two-dimensional shape of the fBOS from a single foot has not been measured in older adults, nor has the area of the fBOS that can support large forces been identified, and it remains unclear if the fBOS can be applied to dynamic tasks such as walking. King et al.^11^ and Tomita et al.^9^ measured the extent of the fBOS in a series of discrete directions under both feet of younger and older adults during the limits of stability test and found that the fBOS is smaller in older adults. Since ground forces were measured under both feet using a single force plate, it is unfortunately not possible to construct an individual foot’s fBOS from their measurements. Hof and Curtze^12^ measured the effective BOS (eBOS) in the sagittal and frontal planes by perturbing standing participants and using a model to infer the margin-of-stability from their resulting movements. While the Hof and Curtze’s methodology^12^ makes it possible to capture the eBOS during a perturbation response, the two directions analysed in their work are insufficient to create a two-dimensional eBOS model of each foot. Melzer et al.^13^ compared the COP area between younger and older adults during quiet standing both with and without a distraction task. Although the results showed a greater sway in older adults, the task did not challenge participants’ balance sufficiently to extract the 2D fBOS. None of these studies have identified a fBOS model that can support large forces and could plausibly be used during large balance corrections. This is an important consideration since only a few parts of the foot are robust enough to support a large fraction of the body’s weight. In addition, previous work has not yet examined to see if the fBOS models extracted from standing tasks can be applied more generally to dynamic tasks such as walking. Finally, the few studies that compare younger and older adults^9,11,13^ make it clear that there are noticeable differences between the two groups, suggesting that an age-specific fBOS model is needed.

In this work, we develop age-specific fBOS models and study how the geometry of these fBOS models vary with age, clinical measures, and correlate with COP length during the stance phase of walking. Our fBOS models of older adults extend existing literature by identifying the two-dimensional shape of the fBOS that can support a large fraction of the body’s weight^10^, and could plausibly be used during a substantial balance correction. To do this work, we first create fBOS models for a group of 38 younger adults, 14 middle-aged adults, and 34 older adults and examine how the fBOS area varies between each group. Second, we analyse how the fBOS area correlates with three common clinical measures that are related to balance problems: frailty level, physical performance, and the fear of falling. Third, we evaluate the correlation between the fBOS length and the COP path length during walking. This final evaluation allows us to see if the fBOS, which is collected from the COP data during a standing task, is related to the COP trajectory during a dynamic task. We hypothesize that the standing fBOS area declines with increasing age^9,11,13^, increasing frailty level^14,15^, decreasing physical performance^16,17^, and increasing fear of falling^18–20^ as these factors are related to increasing occurrence of balance problems and falls. Finally, we hypothesize that a shorter standing fBOS length correlates to a shorter COP path length during walking. To help others build on our work, we have made our scalable fBOS models of the younger, middle-aged, and older adults available online.

## Methods

A group of 34 older adults, 14 middle-aged adults and 38 young adults were included in the study (see Table 1 for details). The frailty level of the older group ranged from very fit to vulnerable on the Clinical Frailty Scale (7 adults of level 1; 13 adults of level 2; 10 adults of level 3 and 4 adults of level 4, where 1 is fit and 4 is vulnerable)^21^. The participants were recruited and assessed at Heidelberg University. Participants were excluded if they had neurological, cardiovascular, metabolic, visual, auditory, mental or psychiatric impairments or injuries that might interfere with the planned task. In addition, participants were excluded if they were unable to independently walk short distances or stand up from a chair. Younger participants had to be between 17-39 years of age, middle-aged participants between 40-65 years of age, and older participants 65 years or older. Older participants were excluded if their frailty level was more than moderately frail on the Clinical Frailty Scale (score ≥6)^21^, or if they were severely cognitively impaired (Mini-Mental State Examination score ≤17)^22,23^. The protocol was approved by the IRB of the medical faculty of Heidelberg University. All experiments were performed in accordance with local guidelines and the Declaration of Helsinki. Participants provided written informed consent.

**Table 1.** Participant characteristics per age group. The mean and standard deviation as well as the [range] are given. The shoe length is derived from the toe and heel markers attached to the shoe. The number of participants having a preference for the right foot (for kicking a ball) versus the left foot (other particpants) are given.

|  | Older adults | Middle-aged adults | Young adults |
| --- | --- | --- | --- |
| Females | 22 / 34 | 6 / 14 | 16 / 38 |
| Age (yr) | 76±6 [66-89] | 50±7 [41-63] | 26±5 [21-39] |
| BMI (kg/m<sup>2</sup>) | 25±4 [17-33] | 24±3 [20-31] | 23±3 [18-30] |
| Height (m) | 1.68±0.09 | 1.79±0.12 | 1.74±0.07 |
| Weight (m) | 72±14 | 77±15 | 70±12 |
| Shoe length (m) | 0.27±0.02 | 0.30±0.02 | 0.27±0.02 |
| Right preference | 20 | 6 | 20 |

### 0.1 Protocol

The fBOS is extracted from the foot position and ground forces as the participant makes the largest possible COP loops while keeping their feet flat on the ground, similar to our previous study in younger adults^10^. Participants wore their own shoes (typically running or similar shoes as requested) that left their ankles uncovered. They stood with feet hip-width apart, toes forward, and with each foot on a force plate. Participants were instructed to move their center-of-mass as much as possible in forward, outwards, backwards, and inward directions in slow, circular movements, while trying not to move their feet or lift their toes or heels. No instructions were given for arm movements or position, or how to best lean in the different directions. If a participant was not effectively moving their COP, they were given examples by the examiner and encouraged to try again.Participants were given as much time as needed and encouraged to make plenty of circular movements. If participants lost their balance the trial was repeated. Participants were not secured in a safety harness as it could affect the assessments, but were spotted by a nearby examiner.

Full body kinematics were recorded at 150 Hz using a motion capture system (Qualisys, type 5+ cameras, Gothenburg, Sweden) and ground reaction forces at 900 Hz using two ground-embedded force plates (Bertec, Columbus, OH, USA). Motion capture markers (14mm) were placed according to the full-body IOR marker model with small adjustments made to the foot markers: on the heel, the medial and lateral ankle epicondyle, the lateral outside of the MTP5 metatarsal, the medial outside of the MTP1 and the front tip of the longest toe (see Figure 2 in^10^). Marker data were labeled in Qualisys.

To examine how the fBOS was related to clinical measures, we assessed the frailty, physical performance, and fear of falling in our older participants. We used the the score on the Clinical Frailty Scale^21^ to assess the frailty level, the total score (and timing on the individual items) on the Short Physical Performance Battery (SPPB)^24^ to assess physical performance, and the score on the short Falls Efficacy Scale International to assess fear of falling (FES-I;^25^). For the SPPB 4m walking test, participants were asked to start on the starting line and walk at a natural pace that feels comfortable. These metrics were not measured from the younger and middle-aged adults because these age groups typically saturate each of these measures.

To examine how the fBOS length relates to the COP path during walking, we extracted the length of the COP trajectory during walking from the kinematics and ground reaction forces collected in the older participants. Participants were asked to walk at a natural pace that feels comfortable^26,27^. Participants walked a distance of roughly 10m for several trials up and down the lab while ground reaction forces were measured with the two ground-embedded force plates and kinematics using the motion capture system. Qualisys software calculated the COP data from the ground reaction forces and moments measured by the force plates. Strides with good full single-foot landing on the force plate were selected by visually inspecting the foot landing in Qualysis as well as checking the force data in MATLAB. Good strides were available for n=33 older adults (n=19 of frailty level 1 or 2, n=10 frailty level 3 and n=4 of frailty level 4). A zero-phase low-pass Butterworth filter with a cut-off frequency of 10Hz was applied to the force data. To exclude data outliers, force and COP data were taken above a threshold of 50N vertical force. The foot-normalised COP path length was calculated by taking the maximum minus the minimum COP points in the anterior-posterior direction in the instructed direction of travel, and dividing these by the foot length calculated from the motion capture markers as above. The medio-lateral direction was not assessed because the COP typically does not traverse the width of the foot during walking.

### 0.2 Functional Base-of-Support

Older adults found the experimental fBOS task particularly challenging: widely circle the COP without moving the feet. As a result, we had to have a detailed set of criteria to identify COP data in which the participant’s foot could be considered to be flat on the ground. Specifically, we selected COP data that had at least 150 consecutive data points (1 second long), supported least 40% of body weight on the respective force plate (which was the main reason for data point exclusion), during which the foot was sufficiently flat on the ground, while excluding the largest 0.1% COP transitions (see^10^ for more details). The resulting force plate data were low-pass filtered using a zero-phase second-order low-pass Butterworth filter with a cutoff frequency of half the sampling rate of the marker data and down-sampled to match the sampling frequency of the marker data.

The fBOS of each participant and each age group were evaluated using the processed COP data. First, we projected the COP data onto a foot-fixed frame that is coincident to the footprint during quiet standing. Next, we computed the fBOS of each foot as the polygon that represents the convex-hull that surrounds the COP data for each foot. We reflected the right foot fBOS polygon medio-laterally (making it equivalent to the left foot) and averaged the convex hull of the left and reflected right feet together. Finally, each participant’s fBOS was normalised by length and width of the foot before the average fBOS was computed for each age group using the arc-length indexed polygon averaging method described in Millard and Sloot^10^. All calculations were performed using custom-written scripts in Matlab (version 2021a, Natick, MA, USA). Our age-specific fBOS models and functions to convert them to different marker sets are available online (see the *Data Availability Statement* for details).

### 0.3 Statistics

The primary outcome is the normalised fBOS area, which is the area of the fBOS divided by the area of the footprint. In this work, the area of the footprint is the area enclosed by the six markers on each foot. Our secondary outcome variables describe the shape of the fBOS, including the inward distance from the heel, front toe, lateral ankle and little toe markers, and the fBOS length and width normalised by the length and width of the foot. Statistical analyses were performed in Matlab (version 2021a, Natick, MA, USA).

First, we examined how the primary and secondary variables differed between the three age groups using a Kruskal-Wallis test, as the data in the individual groups did not appear normally distributed when plotted in histograms. Post hoc comparisons were performed on mean rank values using Matlab’s *multcompare* function. As the age groups have a large age range, we also correlated the normalised fBOS area to age across all data using (non-parametric) Spearman’s correlation.

Second, we investigated how the normalised fBOS area relates to clinical measures of frailty, physical performance, and fear of falling. We performed this analysis only in the group of older adults, as we expected this group to show variability in both the fBOS area size and the clinical measures. Using a Kruskal-Wallis test, we examined the difference in fBOS area between three frailty groups: level 1 & 2 on the Clinical Frailty Scale^21^ (merged into a group of n=20 as they are more difficult to discriminate), level 3 (n=10), and level 4 (n=4). Next, we examined the relation to physical performance by correlating the normalised fBOS area to the Short Physical Performance Battery score (SPPB)^24^, as well as the 4m gait speed and sit-to-stand-to-sit time taken as part of the SPPB, all using Kendall correlations for scores. Finally, we explored the correlation to fear of falling using the score on the Falls Efficacy Scale^25^ using Kendall correlations.

Third, we made a preliminary evaluation of the applicability of the standing fBOS to a dynamic task in older adults. Specifically, we correlated the normalised fBOS length assessed during bipedal standing to the normalised COP path length assessed during walking using available data of 32 older adults (see section 0.1) using a Spearman correlation. We normalised fBOS length and COP path length by the distance between the heel and front toe markers. We only performed this correlation among the older adults because this age group has the largest variability in both fBOS area and walking speed of the three age groups we tested.

## Results

The average fBOS was much smaller than the anatomical outline formed by the foot markers: the median and range were only 22 [9-36]% of the footprint for the young group, 22 [17-29]% for the middle-aged group and 13 [6-28]% for the older group. The fBOS area is longer than it is wide: the maximum fBOS length normalised to foot length was 65 [49-81]% for young, 61 [50-65]% for middle-aged and 53 [22-72]% for older participants. The maximum fBOS width normalised to foot width was 35 [21-62]% for young, 35 [24-39]% for middle-aged and 21 [12-40]% for older participants. Note the large variability between adults within each group.

The fBOS area decreased with age (p*<*0.001). The older group’s median fBOS area was only 58% of the size of the young group (p*<*0.001) and 59% of the size of the middle-aged group (p=0.001; Fig. 1). The individual data points across all age groups also show that the fBOS area correlated to age (Spearman *ρ*=-0.52, p*<*0.0001; Fig. 2). This age-related reduction in fBOS area is direction-dependent: while the inward distances from the lateral ankle (p=0.27) and little toe (p=0.89) do not change, the fBOS edge retreats inwards the foot away from the toes (p=0.0006) and heel (p*<*0.015). Specifically, the inward normalised distance from the toes increases from 21 [10-36]% in young (p=0.0008) and 21 [16-30] % in middle-aged (p=0.025) to 28 [16-43]% of the foot length in older adults. The inward distance from the heel increases from 17 [9-29]% in young (p*<*0.01) to 20 [13-39]% of the foot length in older adults (with 18 [14-28]%, p=0.30, in middle-aged).

**Figure 1.**
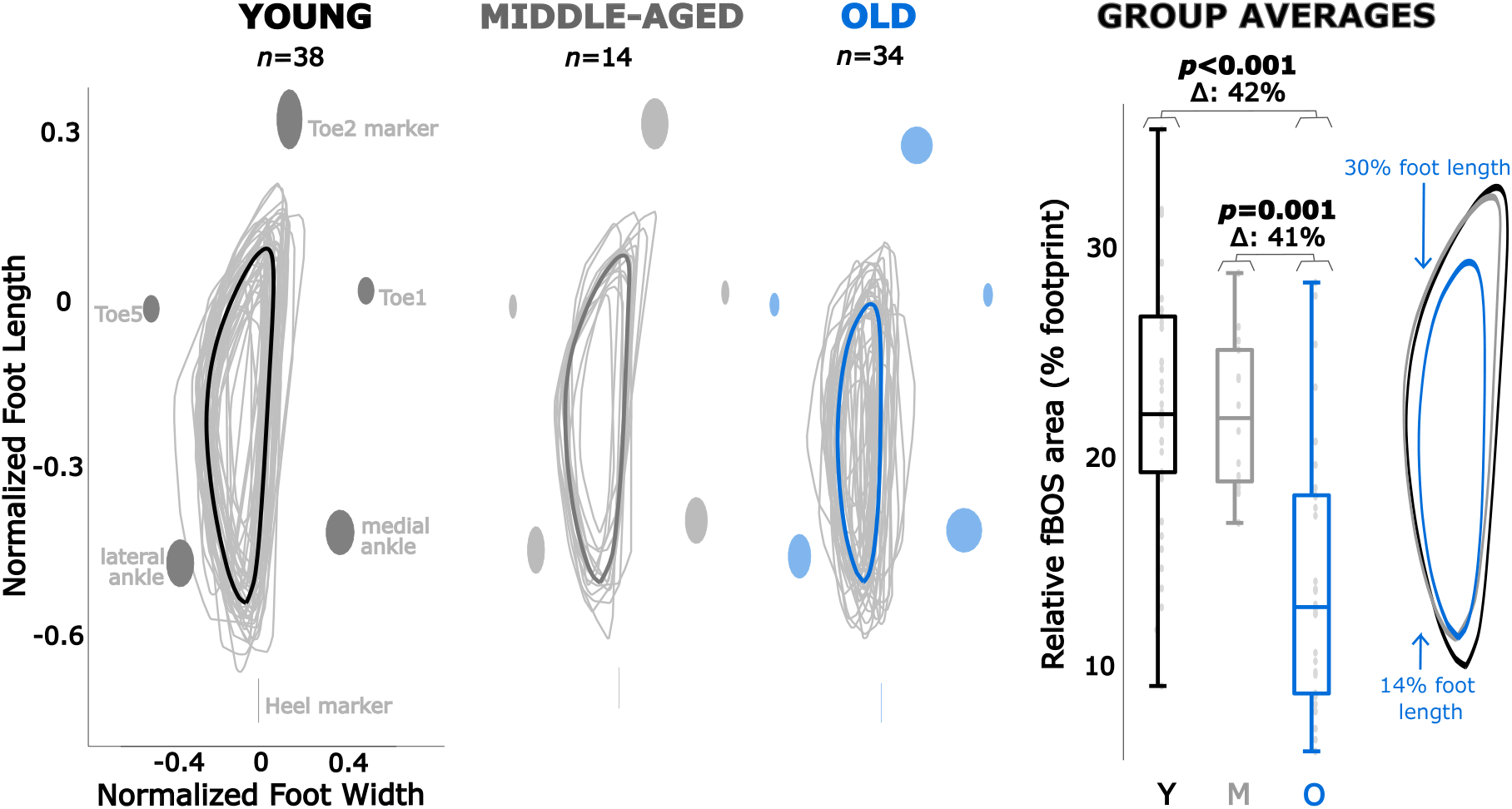
Functional base of support (fBOS) area per age group. Areas are normalised to an individual’s foot length and foot width, with group averages shown in bold. The ellipses show the standard deviation of the marker positions across the group. Note that the heel marker has no mediolateral deviations because the y-axis of the fBOS passes through the heel marker^10^. The boxplots indicate the median and interquartile ranges of the area surface per group, with individual participant’s values indicated with gray dots and outliers with a red cross. The average profiles of the fBOS areas are shown on the right for comparison. Significant differences are indicated with the p-value.

**Figure 2.**
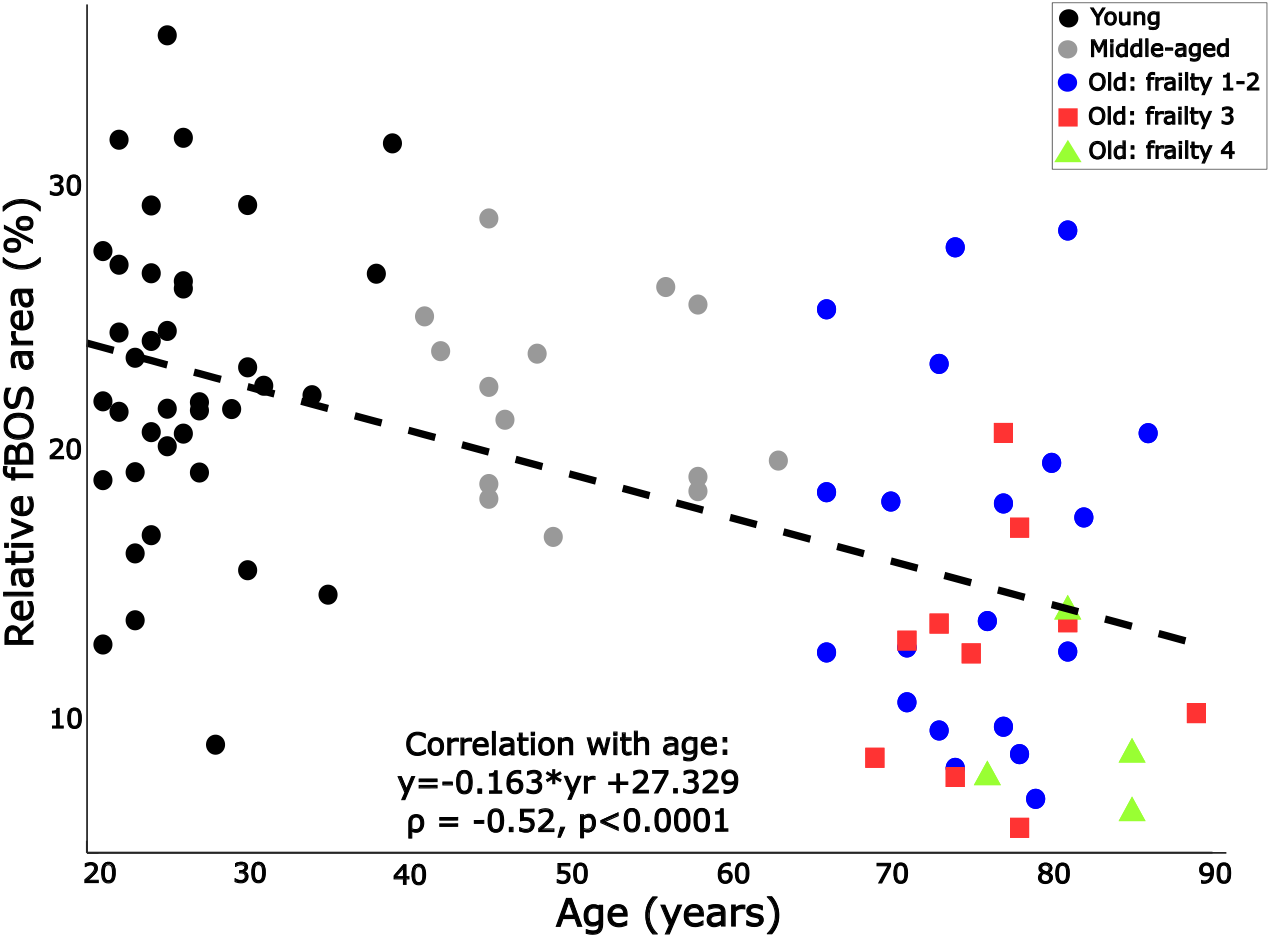
Change in the functional base of support (fBOS) area with biological age. The young adults are shown in black, the middle-aged in grey, and the older adults in colour. While not separately examined here, frailty levels 1 and 2 are shown in red squares, level 3 in blue circles and level 4 in green triangles for context. The relationship was examined using a Spearman correlation (the p-value and *ρ*-value are given, as well as the least-squares first-order linear fit.

In addition to ageing, the size of the fBOS area appears to also be related to frailty and physical performance in the group of older adults. The fBOS area tended to be smaller in older adults with a higher level of frailty (p=0.09; Fig. 3A). The median fBOS area of the pre-frail adults (frailty level 3) was 81% of the non-frail people (level 1 and 2; p=0.54), while the fBOS area of mildly frail adults (level 4) was only 53% of the non-frail people (p=0.10). A decrease in physical performance was weakly correlated with a decrease in fBOS area, both in terms of the total SPPB score (*τ*=0.28, p=0.04; Fig. 3D) as well as the 4m gait speed (*τ*=0.25, p=0.04; Fig. 3B). There was no correlation between fBOS area and SPPB sit-to-stand time (Fig. 3C) or fear of falling (FES; Fig. 3E). Even after the lowest outlier was removed, there was no correlation between fBOS area and SPPB sit-to-stand time.

**Figure 3.**
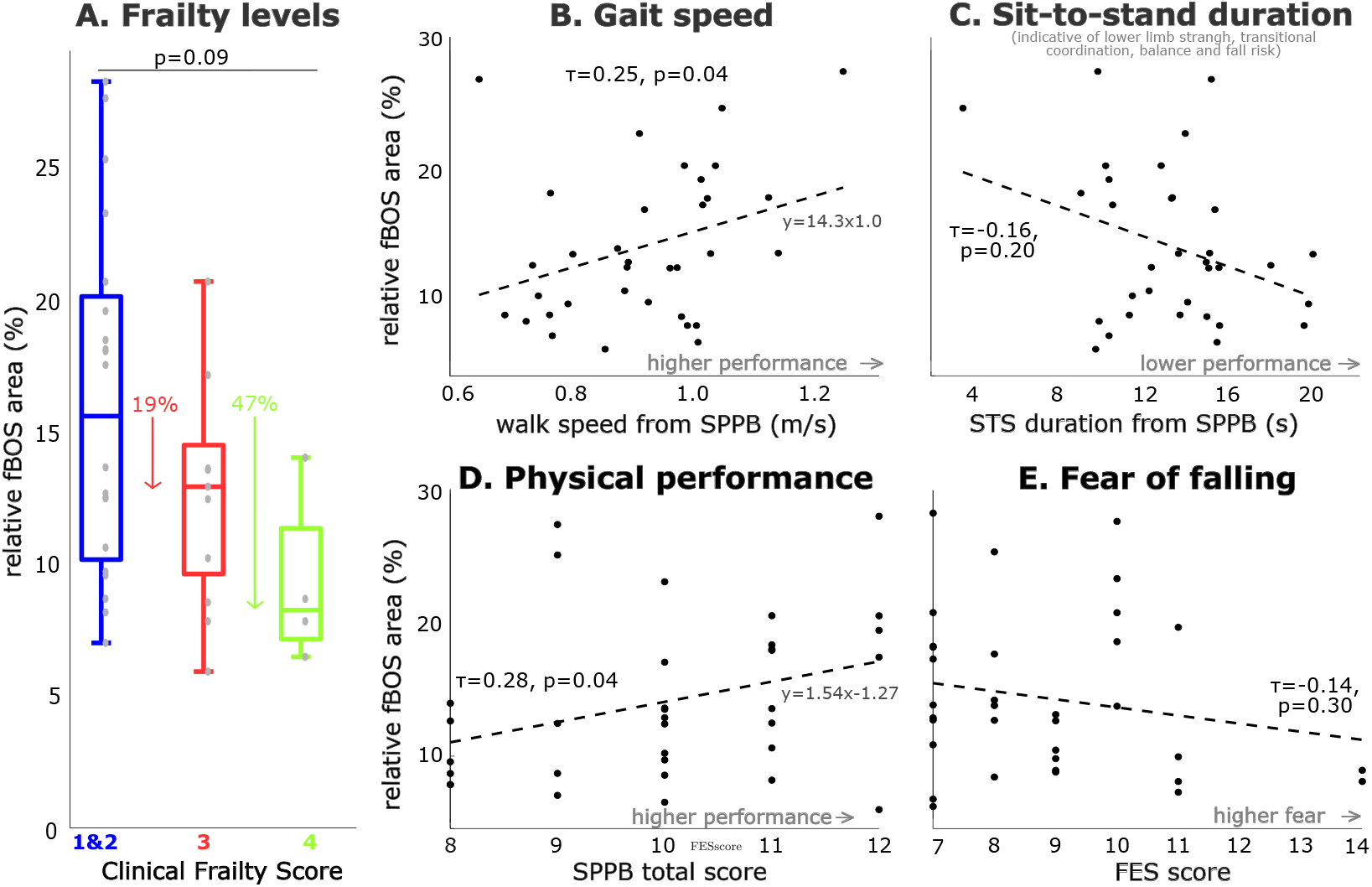
Relation between frailty level, physical performance and fear of falling in older adults versus the functional base of support (fBOS). The relationship of the normalised fBOS area to the foot outline relative to frailty level is shown (A), with frailty based on the Clinical Frailty Scale and with 1 and 2 merged. The percentage decrease for frailty levels 3 and 4 is given relative to the median value of levels of 1 and 2 and includes the level 3 outlier (the reduction is 28% without the outlier). Physical ability was examined as the total Short Physical Performance Scale (SPPB) score (D), the underlying 4m gait speed test (B) and the 5 times sit-to-stand time (C). Fear of falling was established using the Short Falls Efficacy Scale (E). Relationships were examined using Kendall correlations and the least-squares first-order fit to the data points is shown for reference.

Our preliminary evaluation indicates that perhaps the fBOS assessed during standing can be applied to walking. Specifically, we found that the static normalised fBOS length and normalized COP path length measured during walking are correlated (rho=0.49, p=004) among the older adults (Fig. 4). This correlation is across a wide range of static fBOS lengths (22-72% of foot length), and stride speeds (0.82-1.63 m/s) present in the group of 34 older adults.

**Figure 4.**
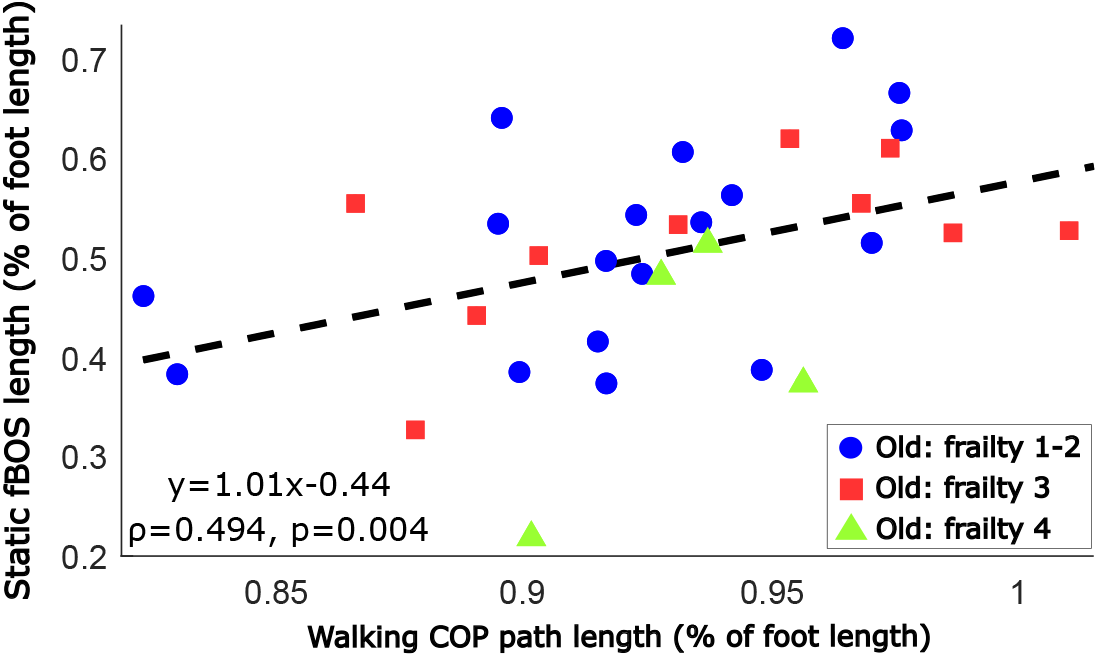
Relation between standing and walking: the standing functional base of support (fBOS) length versus the center of pressure (COP) path length during walking in older adults. The least-squares first-order fit to the data points and the results of the Spearman correlation are given. The colour-coding of the different frailty levels is solely for context.

## Discussion

Injury due to falling is common and on the rise among older adults^1–3^. Model-based balance analysis using the margin-of-stability has the potential to identify people who are at risk of falling before injury occurs, but requires an accurate fBOS model. In this work, we extended our prior fBOS study of younger adults^10^ by making measurements and age-specific geometrical models across a large sample of younger adults, middle-aged adults, and older adults.

Regardless of age, the fBOS is much smaller than the footprint. We found the fBOS area to be only 22% of the area of a single foot across all age groups. This percentage is similar to our prior work with younger adults^10^, but smaller than the 47% reported previously^9^ across two feet. Note that the footprint based on just six foot markers is a slight under-representation of the physical footprint. In addition, we also found that the length of the fBOS in the anterior-posterior direction is 65% of the length of the foot, which is similar to the 50-60% range of prior work^11,28,29^. This emphasizes the importance of using a geometric model of the base of support, such as the fBOS, rather than using motion capture markers to represent the base of support.

We hypothesized that age affects the fBOS area, and this turned out to be partially true: the fBOS of the older adults is the smallest, but there is no difference between middle-aged and younger adults. The fBOS area declined from a mean of 22% of the footprint area in younger adults to a mean of only 13% of the footprint area in older adults, a result that is in line with previous fBOS studies of older adults^9,11,28–33^. Measuring a larger group of middle-aged and older participants is necessary to identify the general age of onset of fBOS decline in future studies. The reduction we found is not uniform but is most pronounced at the toes and heel, with no differences at the sides of the foot. Tomita et al.^9^ also found the largest decrease at the toes, followed by the heel, but they did find a decrease in the lateral direction in older adults, as was found by other studies^9,31,32^. The lack of lateral differences in our results does not seem related to our population since our older adults were similar or even slightly older and frailer to comparable studies^9,31,32^ in the literature.

The reported relationship between age and fBOS area could be dependent on sex. Indeed, females are more prone to falling than males^34,35^. This could be caused by sex-related differences in kinematic and kinetic gait parameters^36–38^, postural control^39–41^, or the reported lower balance performance, gait stability and foot clearance found in females^34,40,42–44^. A large cohort study did not find differences between males and females in margin-of-stability^45^. However, Meltzer et al.^46^ found that females tend to have a smaller foot-normalised anterior-posterior limit of stability. An additional analysis did not reveal any differences between females and males in fBOS area across all participants (Wilcoxon rank sum test: p=0.44), for the young group (p=0.75) or the old group (p=0.60). Thus, there is no indication that sex modified the decline in fBOS with ageing that we found.

We found considerable variability between participants. As seen for the discrete limits of stability^9,11^, some older participants have fBOS areas that are larger than those of some younger participants. Even for the young, the fBOS areas range from 9 to 36%, with the middle-aged and older adults showing a similarly large spread in values (between 3-29%). As the task required to measure the fBOS is challenging, these differences may arise due to differences in strength^18,46,47^, coordination^30–32^, sensory acuity^48–50^, or perhaps due to the structure of the foot (including stiffness of the feet and ankle joint) rather than a difference in balance ability.

Our hypothesis that fBOS area would correlate with clinical measures is directionally correct, though not all correlations are statistically significant. The fBOS tended to decrease with increasing frailty level. This relation might become more pronounced when measuring a larger group of more frail older adults. The fBOS area decreased with both the clinical SPPB test of physical performance and comfortable walking speed. Particularly, the relation with walking speed is interesting, as walking speed is considered a functional vital sign and predictive of mental health, disability, falls, physical activity and more^51,52^. We did not find a correlation with fear of falling using the FES scale in contrast to Binda et al.^18^, who found a correlation between anterior-posterior limits and scores on the Activities-specific Balance Confidence (ABC) scale. It is worth noting that Binda et al.^18^ specifically recruited people who had a fear of falling; as we did not our participants did not report much fear of falling. A relation between fBOS anterior-posterior limits versus fall history and balance recovery has been reported^28,53,54^. Unfortunately, we could not investigate such relations using the fBOS area as only a few individuals were fallers or had a low balance score on the SPPB. While these relations generally align with those known for balance, there is much variability and thus biological age seems currently the most promising factor to account for in fBOS to improve balance analysis accuracy.

Since we have found that the size of the fBOS correlates with the length of the COP trajectory during walking, it is possible that the fBOS could also be applied during dynamic movement. This is despite people not necessarily moving towards the boundaries of their fBOS during steady-state, unperturbed walking. To rule out that this relationship is actually caused by both the fBOS and the COP being related to walking speed (we already showed this relation for the fBOS in Fig. 3B), we evaluated the relation between COP and walking speed. We did not find a correlation between the COP path length and walking speed calculated for the same strides (Kendall Tau: 0.057, p-value: 0.66). Future work should focus on establishing the validity of using the fBOS to predict the area that bounds COP trajectories during dynamic movement.

Our results may be limited by the task we used to measure the fBOS, group size, and the footwear worn by participants. First, we can never be sure participants reached their boundaries during the whole circular motion, though compensatory arm movements made it clear that many found the task challenging. As a result, differences in motivation, fear of falling, and adherence to instructions could have enlarged the variability in fBOS area between participants. Next, while our group size should be sufficient for the main comparisons between age groups, more older and particularly frail adults would have improved the power of the correlation and secondary analyses. It should be noted that one participant (72 yrs, frailty level 2) was not included in the population due to an abnormally large fBOS area that we think is related to inappropriate thresholds for foot movement for this participant (see Millard and Sloot^10^). Finally, we did not control for the stiffness of the shoe or the stiffness of the foot. Previous studies examining the limits of stability measured participants barefoot^9,11,46^. We did not find any systematic differences between barefoot and shod fBOS areas in our prior study of 27 younger adults^10^. We compared data available for 8 middle-aged and 3 older adults (see the Supplemental Data file) and found no difference between shod (21.20% [8.25%]) and barefoot (20.06% [9.26%]) conditions (p signrank = 0.90). Since we did not control for footwear, nor the stiffness of participants’ feet (flat foot vs. high arch), it is possible that these factors affected our results^55,56^. Future studies should examine the effect of different types of footwear and foot stiffness on the fBOS.

In constructing the planar fBOS model we have presented here we have had to make a number of practical decisions along the way. We have chosen to assess the shape of the fBOS of a single foot across each age group to be able to apply our model to walking, including the single-stance phases. We describe the fBOS as a two-dimensional polygon^10^ because this shape makes it possible to accurately assess the margin-of-stability in any direction, rather than being limited to the specific measurement directions of prior work. We averaged over the left and right foot, as we previously did not find an effect of dominant leg in young persons^10^. As some participants struggled to indicate their dominant leg, we performed furfther analysis to see if focusing on the leg with the smallest fBOS area would have changed the results. We found a similar correlation to age for the smallest fBOS area compared with the average over the left and right foot (*ρ*=0.47, p*<*0.001).

Our measurements of younger, middle-aged, and older adults make it possible to scale an age-specific fBOS template to a participant to evaluate the margin-of-stability more accurately than would be possible otherwise. Our fBOS models offer far better accuracy than using motion capture markers to directly define the BOS. In younger adults, our fBOS model will improve the accuracy of the margin-of-stability by 6.5 cm from the toes, 5.2 cm from the heel, and 2.0 cm from the side (at the height of TMP5), for an average-sized foot of 30 cm by 10 cm. When the age-specific fBOS model is applied to older adults the improvements in accuracy increase to 8.3 cm at the toes and 6.5 cm at the heel. These improvements in accuracy are large in comparison to the size of the foot and will make it easier to detect people who have balance problems before injury occurs. The provided age-specific models (applicable to the IOR^57^ or PlugInGait^58,59^ marker sets with code to convert to other marker models) thus offer a generally applicable definition for the fBOS which can be applied to foot marker data recorded during different movements to analyse balance.

## Conclusion

We measured the functional base of support across younger, middle-aged and older adults and found that this fBOS area decreases with age. This relation was stronger than the decline in fBOS area we found with reduced physical ability and the trend of decline with frailty level. The fBOS measured during standing might be representative to the fBOS boundaries during walking, as it correlates to the length of the COP path during stance. So that others can extend our work and apply it to their own research, we have made the age-specific fBOS models and example data available online (see the *Data Availability Statement* for details).

## Acknowledgments

The authors would like to thank the participants for their time and contribution. We would like to acknowledge the support of our student assistant Joel Grathwohl for his help with data collection and processing.

## Author Contributions

LHS and MM designed the experiment; LHS and TG were responsible for participant recruitment, data collection and data processing; MM and LHS created the fBOS code and analysis pipeline respectively; LHS and MM analyzed and interpreted the data; LHS and MM wrote the manuscript and generated the figures; TG, KM and MM, provided critical feedback on the data interpretation and manuscript. KM and LHS obtained funding for the work.

## Data Availability Statement

The outcome variables for all individuals that are used for statistical analysis are made available in the supplementary material file *SupMat_individuals_datavalues_v2*.*xlsx*. The raw data files are available from the corresponding authors on reasonable request. The fBOS models, functions to calculate the fBOS model from data, convert fBOS models to different marker sets, and to average fBOS convex hulls, as well as example data to use these functions, are shared at GitHub: https://github.com/mjhmilla/functionalBaseOfSupport.

## License Statement

The contents of this publication *The size of the functional base of support decreases with age* © 2025 by Lizeth H. Sloot, Thomas Gerhardy, Katja Mombaur, and Matthew Millard are licensed under CC BY 4.0. To view a copy of this license, visit https://creativecommons.org/licenses/by/4.0/

## Additional Information

The authors declare that the research was conducted in the absence of any commercial or financial relationships that could be construed as a potential conflict of interest.

